# Nascent extracellular matrix converts biomaterial cues into cell fate decisions

**DOI:** 10.64898/2026.01.10.698826

**Authors:** J.Y. Liu, E. M. Plaster, M. Fan, D. W. Ahmed, A. Roy, P. Duran, P. Panovich, A. S. Piotrowski-Daspit, C. A. Aguilar, M. L. Killian, C. Loebel

**Affiliations:** Department of Bioengineering, University of Pennsylvania, USA; Center for Precision Engineering for Health (CPE4H), University of Pennsylvania, USA; Department of Biomedical Engineering, University of Michigan, USA; Department of Materials Science & Engineering, University of Michigan, USA; Internal Medicine - Pulmonary & Critical Care Medicine Division, University of Michigan, USA; Department of Orthopaedic Surgery, University of Michigan, USA

## Abstract

Hydrogels serve as powerful models for investigating cell-extracellular matrix (ECM) interactions. While chemical modifications are routinely used to tune hydrogel properties, it remains unclear whether these modifications mediate cell fate. Previous work has shown that cells deposit newly synthesized (nascent) ECM at the cell-hydrogel interface. Here, we demonstrate that this nascent ECM interface regulates how cells interpret chemical modifications. Using hydrogels with varied chemical modifications, we isolated the effects of chemical modification on nascent ECM and cell fate. Nascent ECM deposition increased as a function of hydrogel modification and with distinct matrisome compositions. While low modification hydrogels promoted cell differentiation, high modifications increased cell proliferation. Perturbing cell-nascent ECM interactions reversed this cell fate. Our findings reveal that nascent ECM regulates cell fate by converting hydrogel cues into signals that control cell fate. This tri-directional interplay among hydrogel chemical modifications, nascent ECM, and cell fate reframes how we interpret cell-hydrogel interactions.

## Introduction

The extracellular matrix (ECM), an organized assembly of structural and signaling molecules, integrates mechanical and biochemical signals to regulate cellular responses in tissue homeostasis and disease progression^1^. Engineered hydrogels have emerged as powerful tools to recapitulate key features of the ECM, including the mechanical and biochemical signals that cells are exposed to. Building upon the concept of bi-directional cell-ECM signaling (Mina Bissel, 1982^2,3^), these hydrogels have been instrumental in investigating how the ECM instructs cell function and how in turn cells remodel the ECM. However, recent work has challenged this bi-directional framework by demonstrating that cells upon embedding into hydrogels rapidly deposit a layer of newly synthesized (nascent) ECM (nECM) at the cell-hydrogel interface^4–7^. Importantly, cellular interactions with this nECM have been shown to regulate mechanosensing^8,9^ that directs downstream cell function such as cell spreading and differentiation^4,10^. Moreover, nECM accumulation has been reported to support the formation and growth of three-dimensional multicellular structures including spheroids and organoids *in vitro*^10,11^. While hydrogel mechanical properties are known to regulate the thickness of deposited nECM^8,10,12^, how the hydrogel polymer backbone itself contributes to nECM deposition and cell function remains unexplored. Thus, there is a critical gap in our fundamental knowledge on the role of hydrogel-nECM interactions in regulating cell function and fate.

The fabrication of hydrogels often requires chemical modifications of the polymer backbone (e.g., methacrylates/acrylates, norbornenes, or vinyl-sulfones) to form crosslinks and provide handles for biochemical signals (e.g., RGD (Arginine-Glycine-Aspartic acid), peptides or full-length proteins)^13,14^. While recent work highlighted that RGD ligand presentation reduces nECM deposition by spheroids in methacrylated hyaluronic acid hydrogels^6^, whether the methacrylates themselves alter nECM deposition remains unknown. Interestingly, a previous study suggested that increasing modifications of hyaluronic acid hydrogels impair chondrogenic differentiation of mesenchymal stromal cells^15^. Yet, how the chemical modifications of the hydrogel polymer backbone alter nECM deposition, composition, and its functional properties are unknown.

To address this knowledge gap, we synthesized hyaluronic acid polymers with varying degree of norbornene modifications to fabricate hydrogels with matched crosslinker concentration and mechanical properties. Building upon previous studies^9,16^, we used metabolic labeling to characterize the spatiotemporal evolution and composition of cell-deposited nECM as a function of chemical modifications. In addition, we performed perturbation studies to dissect the specific contributions of cell-hydrogel and cell-nECM interactions in guiding cell fate decisions. Our findings show that hydrogel modification, independent of mechanical properties, regulated nECM deposition and cell fate. This work challenges the traditional bi-directional interaction between cells and engineered hydrogels and instead provides evidence for a tri-directional interplay among hydrogels, nECM, and cells within 3D hydrogel culture.

## Results and discussion

### Hydrogel modifications direct nECM deposition

To probe how hydrogel modifications direct nECM deposition, we engineered norbornene-modified hyaluronic acid (NorHA) hydrogels with ‘low’ (∼10%), ‘mid’ (∼23%), and ‘high’ (∼43%) degree of norbornene modification (Figure 1a). NorHA hydrogels were then crosslinked via thiol-ene reaction using dithiothreitol (DTT) and the mechanical properties were measured via compression testing. Using 2 wt% NorHA, increasing the DTT concentration increased the Young’s modulus across groups (Figure 1b). However, the Young’s moduli saturated at different DTT concentrations depending on the degree of norbornene modification. For ‘low’ modification hydrogels, saturation occurred at approximately 3.25 mM DTT, whereas ‘mid’ modification saturated at 6.5 mM DTT and ‘high’ modification at 9.75 mM DTT. This is expected due to the differences in the amount of norbornene available for crosslinking^16^. In contrast, hydrogels crosslinked with 0.65 and 1.3 mM DTT exhibited comparable Young’s moduli. Thus, to investigate the contributions of hydrogel modification independent of mechanical properties, we selected 1.3 mM DTT to obtain 5 kPa NorHA hydrogels with ‘low’, ‘mid’, and ‘high’ norbornene modification. Next, we embedded juvenile bovine chondrocytes in 5 kPa ‘low’, ‘mid’ and ‘high’ NorHA hydrogels and cultured the constructs in methionine-free, L-azidohomoalanine (AHA) containing media, which enabled the visualization and analysis of newly synthesized proteins, using copper-free strain-promoted azide-alkyne cycloaddition with a DBCO (Dibenzocyclooctyne)-fluorophore^9^ (Figure 1c). Within one day of culture, DBCO staining showed a thin and discontinuous nECM layer around cells within all NorHA hydrogels, which then continuously increased in thickness over 7-day of culture (Figure 1d). Within ‘low’ hydrogels, quantification of nECM thickness showed an almost two-fold increase from 0.68 ± 0.54 µm (Day 1) to 1.68 ± 0.74 µm at day 4 but no further increase till day 7 (1.58 ± 0.67 µm, Figure 1e), indicating that nECM deposition plateaued within the first few days of culture. In contrast, nECM thickness of cells cultured within ‘mid’ NorHA hydrogels continuously increased from 0.74 ± 0.55 µm (day 1) to 1.30 ± 0.78 µm (day 4), and 1.59 ± 0.85 µm (day 7), suggesting prolonged nECM deposition when compared to ‘low’ hydrogels (Figure 1f). In ‘high’ hydrogels, nECM thickness was initially comparable to ‘low’ and ‘mid’ hydrogels at Day 1 (0.54 ± 0.31µm) but showed even more significant increases between the time points (Figure 1g-i, day 4: 1.66 ± 1.01 µm, day 7: 2.21 ± 1.26 µm). When comparing nECM thickness across groups, at day 7 we observed a 1.5-fold increase in nECM thickness between cells cultured in the ‘high’ hydrogels when compared to both ‘low’ and ‘mid’ hydrogels, with little differences at day 1 and day 4 (Supplementary Figure 1a-c). Notably, assessing nECM thickness at the single cell level showed little heterogeneity at day 1 whereas by day 7, cells in ‘high’ hydrogels showed a strong increase in heterogeneity as well (Supplementary Figure 1d-f). These data show that hydrogel modifications further regulate the heterogeneity of nECM accumulation.

Taken together, these findings demonstrate that hydrogel modifications guide nECM deposition and heterogeneity throughout the culture time. More specifically, ‘low’ hydrogels promote faster but overall lower nECM deposition compared to ‘high’ hydrogels with sustained and increased nECM accumulation over time.

**Figure 1.**
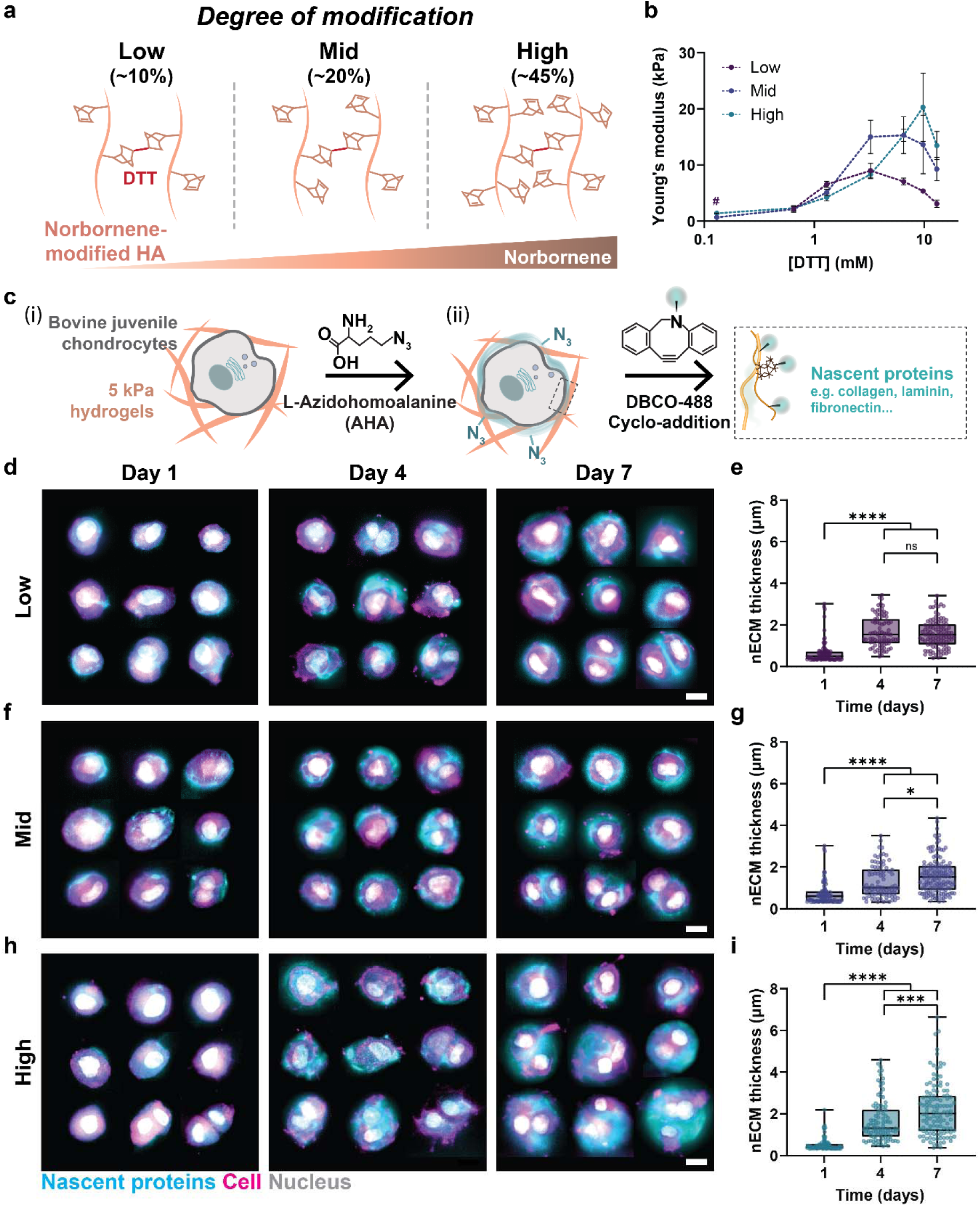
Hydrogel modifications direct nECM deposition and accumulation. **a.** Schematic illustrating the design of hydrogels with varying degrees of norbornene chemical modification (low, mid, high). **b.** Compressive modulus of hydrogels as a function of crosslinker concentration, measured via unconfined compression testing (*n* = 3-4 hydrogels per group, *N* = 3, Day 1). # Not measurable due to lack of structural integrity (low, 0.13 µM). **c.** Schematic of NorHA hydrogel embedded bovine juvenile chondrocytes at 5x10^6^/mL seeding density (i), cultured in media supplemented with azidohomoalanine (AHA), prior to fluorophore-conjugated DBCO click chemistry to visualize nascent extracellular matrix (nECM, iii): **d.** Representative fluorescent images and **e.** quantification of average nECM thickness of chondrocytes cultured in low-modification hydrogels at day 1, day 4, and day 7 (scale bar = 10 μm, day 1 *n* = 85 cells, *N* = 2; day 4 *n* = 82 cells, *N* = 2; day 7 *n* = 101 cells, *N* = 3). **f.** Representative fluorescent images and **g.** quantification of average nECM thickness of chondrocytes cultured in mid-modification hydrogels at day 1, day 4, and day 7 (scale bar = 10 μm, day 1 *n* = 86 cells, *N* = 2; day 4 *n* = 86 cells, *N* = 2; day 7 *n* = 136 cells, *N* = 3). **h.** Representative fluorescent images and **i.** quantification of average nECM thickness of chondrocytes cultured in high-modification hydrogels at, day 1, day 4, and day 7 (scale bar = 10 μm, day 1 *n* = 89 cells, N = 2; day 4 *n* = 102 cells, *N* = 2; day 7 *n* = 122 cells, *N* = 3). **a-i.** *N* = number of independent experiments, error bar = standard deviation, ****p < 0.0001, ***p < 0.001, *p < 0.05, ns: not significant by one-way ANOVA with Tukey’s multiple comparisons test.

### Hydrogel modifications regulate cell proliferation, morphology, and differentiation

After having shown that hydrogel modifications direct the amount and heterogeneity of deposited nECM, we next sought to investigate cellular responses. Hoechst staining at day 7 revealed a higher number of cells with divided nuclei in ‘high’ hydrogels when compared to cells cultured in ‘low’ and ‘mid’ hydrogels (Figure 2a). Quantification of the percentage of divided cell nuclei per total number of cells as a function of hydrogel modification showed a continuous increase over the 7 days of culture, reaching up to 63.47 ± 10.21% of cells with divided nuclei in ‘high’ hydrogels (Figure 2b). In contrast, cells in the ‘low’ and ‘mid’ hydrogels showed an initial increase in dividing nuclei, reaching 33.81 ± 8.22% (low) and 28.00 ± 7.31% (mid) at day 4. After this point, no further increase was observed. When comparing the number of divided nuclei per cell at day 7, we measured an almost 3-fold increase between ‘low’ and ‘high’ hydrogels (Figure 2c), suggesting that hydrogel modifications have a direct effect on cell division. Increased cell proliferation was further confirmed by 5-ethynyl-2’-deoxyuridine (EdU) incorporation over 7 days of culture (Figure 2d). While we observed proliferating cells in all hydrogels, the percentage of EdU-positive cells showed up to a 2.5-fold increase in ‘high’ hydrogels (80.4 ± 8.0%) when compared to ‘low’ hydrogels (31.5 ± 17.4%, Figure 2e). These results suggest that an increase in hydrogel modifications increases cell proliferation. Interestingly, the higher proliferative capacity correlated with higher cell aspect ratios on day 7 (Figure 2f), suggesting that hydrogel modifications may also direct cell elongation during proliferation.

We next sought to determine whether hydrogel modifications alter downstream chondrogenic differentiation, focusing on transcription factor Sox9 (SRY-Box Transcription Factor 9), its downstream transcriptional targets *COL2A1* (Collagen type II), *ACAN* (Aggrecan) and indirect suppression of *COL1A1* (Collagen type I), *VCAN* (Versican) in ‘low’ and ‘high’ hydrogels^17,18^ (Figure 2g). At day 7, immunofluorescence showed increased Sox9 protein nuclear staining and an increase in Sox9 nuclear-to-cytoplasmic ratios for cells cultured in ‘low’ hydrogels compared to ‘high’ hydrogels (Figure 2h). Gene expression of Sox9 was similarly increased in ‘low’ hydrogels compared to ‘high’ hydrogels (Supplementary Figure 2). Cells in ‘low’ hydrogels showed a 3.5-fold increase in COL2A1/COL1A1 and a 1.4-fold increase in the ACAN/VCAN ratios when compared to ‘high’ hydrogels (Figure 2i), reflecting enhanced chondrogenic differentiation^19^. Together, these results demonstrate that ‘low’ hydrogels show a higher tendency to maintain chondrogenic phenotype, compared to ‘high’ hydrogels that promote cell proliferation.

**Figure 2.**
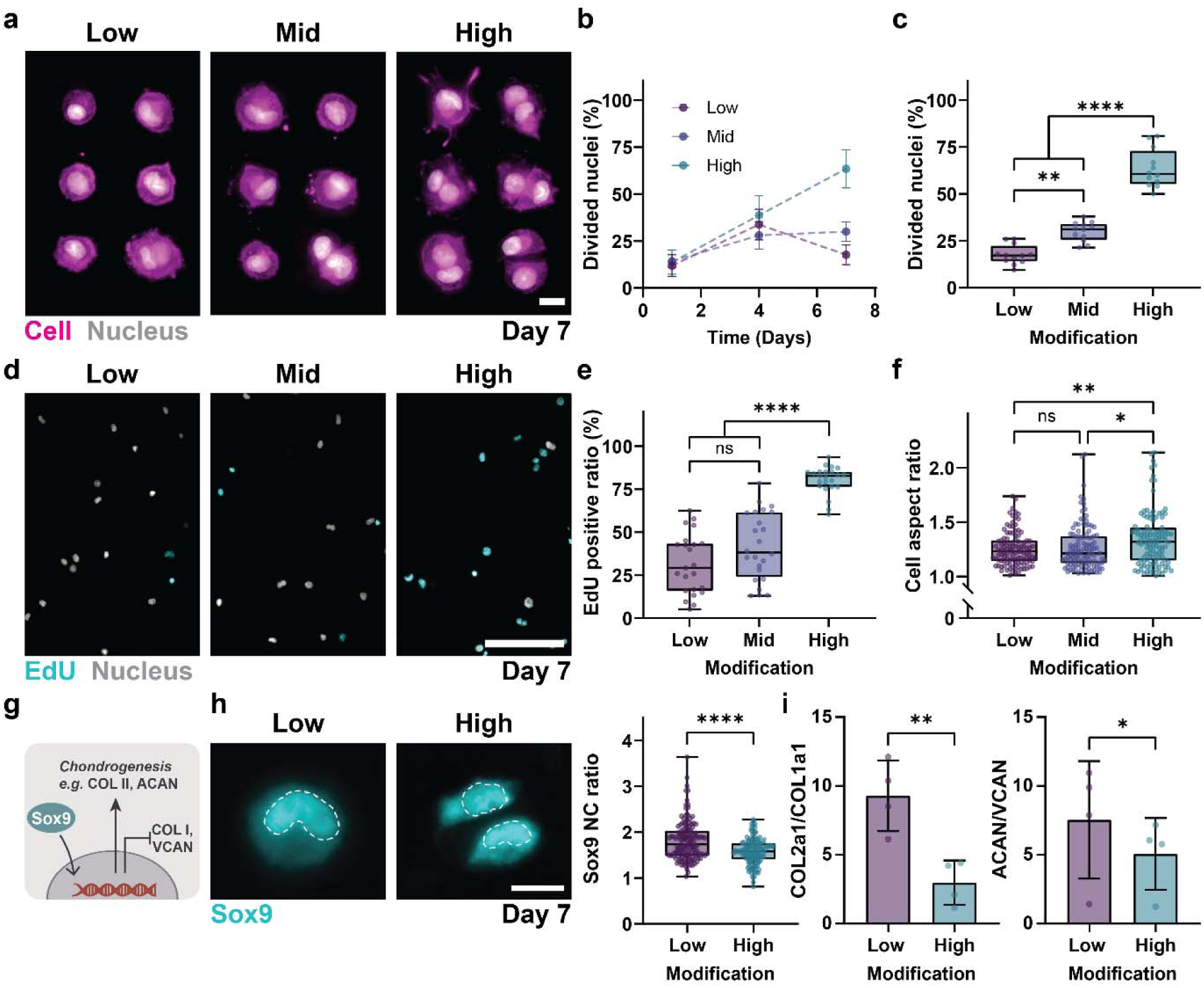
Hydrogel modifications regulate cell fate. **a.** Representative fluorescent images of nuclei and cell membrane of chondrocytes cultured in low, mid, and high modification hydrogels at day 7 (scale bars: 10 μm.) **b.** Quantification of divided nuclei of chondrocytes culture in low, mid and high modification hydrogels at day1, 4 and 7 (low: day 1 *n* = 12 regions of interest (ROIs), *N* = 2; day 4 *n* = 9 ROIs, *N* = 2; day 7 *n* = 12 ROIs, *N* = 3; mid: day 1 *n* = 8 ROIs, *N* = 2; day 4 *n* = 8 ROIs, *N* = 2; day 7 *n* = 12 ROIs, *N* = 3; high: day 1 *n* = 10 ROIs, *N* = 2; day 4 *n* = 9 ROIs, *N* = 2; day 7 *n* = 12 ROIs, *N* = 3) **c.** Quantification of divided nuclei of chondrocytes culture in low, mid and high modification hydrogels at day 7 ( low: *n* = 12 ROIs, *N* = 3; mid: *n* = 12 ROIs, *N* = 3; and high: *n* = 12 ROIs, *N* = 3). **d.** Representative fluorescent images of incorporation of 5-ethynyl-2’-deoxyuridine (EdU) into chondrocytes cultured in low, mid and high modification hydrogels at day 7 (scale bar: 100 μm). **e.** Quantification of EdU incorporation into chondrocytes cultured in low, mid and high modification hydrogels at day 7 (scale bar: 100 μm, low: *n* = 23 ROIs, *N* = 4; mid: *n* = 24 ROIs, *N* = 4; and high: *n* = 23 ROIs, *N* = 4**). f.** Quantification of chondrocyte cell aspect ratio at day 7 (low: *n* = 108 cells, *N* = 3; mid: *n* = 112 cells, *N* = 3; and high: *n* = 111 cells, *N* = 3). **g.** Schematic showing the downstream effects of Sox9 translocation into the nucleus, including upregulation of collagen type IIA1, aggrecan (ACAN) and Sox9, and the downregulation of collagen type IA1, versican (VCAN) and THY-1 **h.** Representative immunofluorescent images and quantification of Sox9 nuclear translocation calculated by the nucleus-to-cytoplasm (NC) ratio at day 7 (scale bar: 10 μm, low: *n* = 139 cells, *N* = 3; and high: *n* = 124 cells, *N* = 3). **i.** Ratio of COL2A1/COL1A1 and ACAN/VCAN gene expression at day 7, measured by qPCR and normalized to S18 housekeeping gene. Gene expression ratios were calculated as 2^(-(ΔCt₁ - ΔCt₂))^, where ΔCt₁ and ΔCt₂ represent Ct values of each gene normalized to S18. (*N* = 4) **a-i.** *N* = number of independent experiments, error bar = standard deviation, ****p < 0.0001, **p < 0.01, *p < 0.05, ns: not significant by one-way ANOVA with Tukey’s multiple comparisons test.

### Initial hydrogel cues preserve cell phenotype

Given that cell fate was altered in response to hydrogel modification, we next investigated how the initial cell-hydrogel interactions act as upstream determinants of cell fate. It has been well established that cells recognize hyaluronic acid via the cell surface receptor CD44, which has been implicated in chondrogenic differentiation^20–22^. Previous studies have shown that cells express CD44 within an hour of being in contact with HA hydrogels^23^. However, the contributions of these early cell-hydrogel interactions to cell proliferation as a function of hydrogel modification have not been well established. Thus, to test how CD44-hydrogel interactions direct chondrogenesis, we used a function-perturbing CD44 antibody. The antibody was added for an hour prior to embedding and throughout the first three days of culture to block CD44 during nECM deposition (Figure 3a). Within ‘low’ hydrogels, blocking CD44 signaling induced an almost 2-fold increase in the number of EdU-positive cells (48.80 ± 8.09%) at day 7, which is similar to the number of EdU-positive cells in ‘high’ hydrogels without CD44 inhibition (Ctrl, Figure 3b). Note that CD44 inhibition had little influence on EdU incorporation in ‘high’ hydrogels (Supplementary Figure 3a), indicating that the initial interaction with ‘high’ hydrogels prevents cell proliferation. In addition, blocking CD44 signaling significantly decreased Sox9 nuclear staining, resulting in a reduction of the Sox9 nuclear-to-cytoplasmic ratio from 1.79 ± 0.41 (Ctrl) to 1.55 ± 0.29 (CD44i), which is similar to cells in ‘high’ hydrogels without CD44 inhibition (Figure 3c). Blocking CD44 interactions in ‘high’ hydrogels did not induce detectable changes in Sox9 staining and nuclear-to-cytoplasmic ratios (Supplementary Figure 3b), These findings indicate that CD44-mediated interactions with the hydrogel backbone are critical towards chondrogenic maintenance.

**Figure 3.**
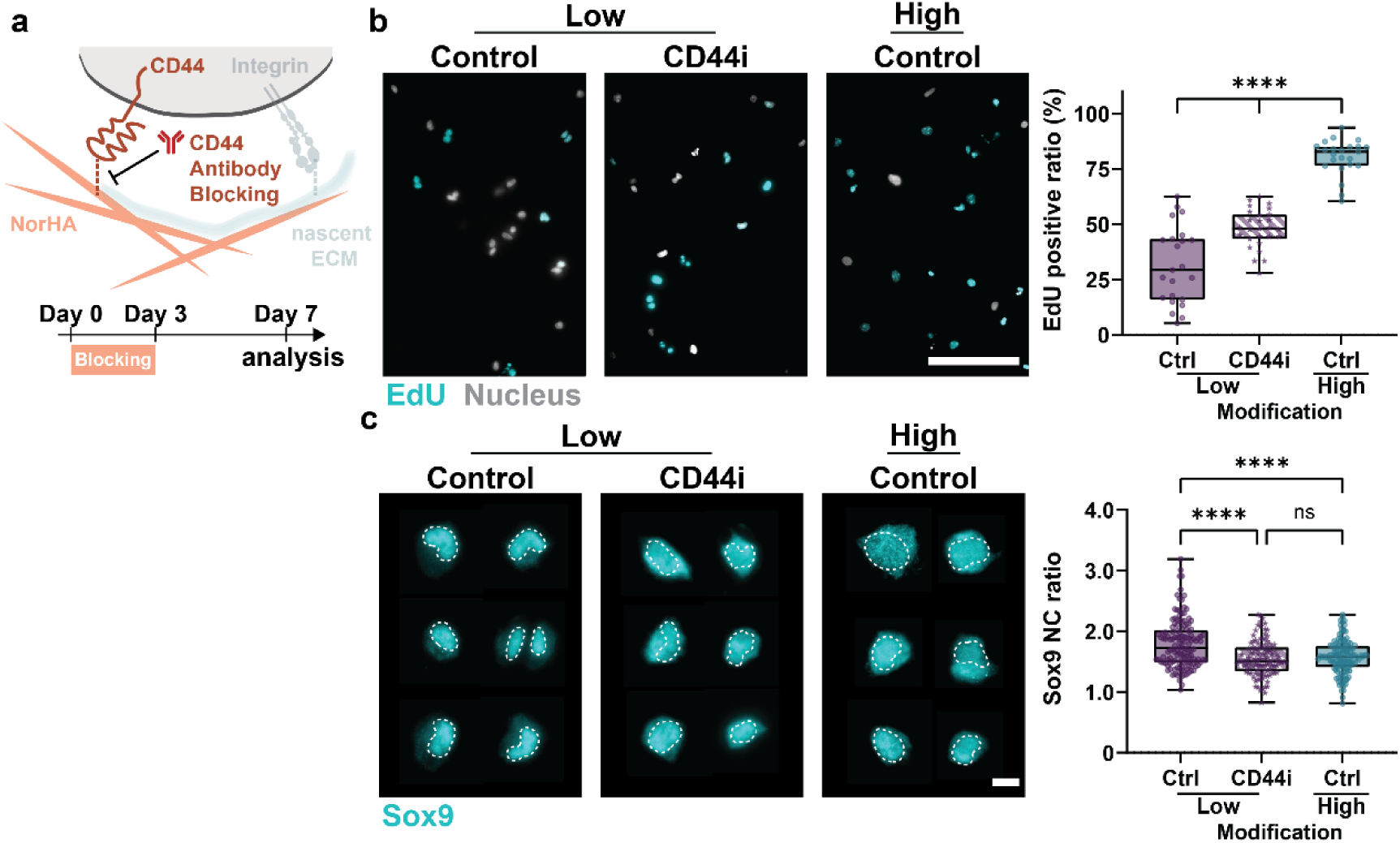
Cellular interactions with the hydrogel backbone are required for maintaining cell fate. **a.** Timeline and schematic showing the blocking of cell adhesion to hyaluronic acid with CD44 function-perturbing antibody for 1 hour pre-embedding and for 3 consecutive days while maintaining adhesion to nECM. **b.** Representative fluorescent images and quantification of the incorporation of EdU in chondrocytes cultured without (Ctrl) and with CD44 inhibition (CD44i) in ‘low’ modification hydrogels at day 7. (scale bar = 100μm, low-Ctrl: *n* = 23 ROIs, *N* = 3; low-CD44: *n* = 27 ROIs, *N* = 3; high-Ctrl: *n* = 23 ROIs, *N* = 3) **c.** Representative fluorescent images (dashed line outlines nuclei) and quantification of Sox9 nucleus-to-cytoplasm (NC) ratio of chondrocytes cultured without (Ctrl) and with CD44 inhibition (CD44i) in low modification hydrogels at day 7. (scale bar = 10μm, low Ctrl: *n* = 138 cells, *N* = 4; low CD44i: *n* = 110 cells; *N* = 4, high-Ctrl: *n* =124 cells; *N* = 4) **a-c.** *N* = number of independent experiments, error bar = standard deviation, ****p < 0.0001, ns: not significant by one-way ANOVA with Tukey’s multiple comparisons test.

### Specific nECM components are spatially heterogeneous at the cell-hydrogel interface

Having established that hydrogel modifications modulate nECM deposition and cell fate, we next sought to examine the compositional and spatial architecture of specific nECM components. Thus, in addition to nECM labeling as a global representation of all newly secreted ECM proteins, we performed immunostaining of well-studied ECM markers at day 7. Co-staining for total collagen by collagen hybridization peptide after thermal denaturation showed high structural similarity with nECM labeling, suggesting that the deposited nECM is rich in collagens (Figure 4a). Importantly, both total collagen and nECM staining showed a continuous pericellular shell but variable intensities across the depth of the cell-hydrogel interface. Overexposure of both stains revealed fibrillar ECM that extended much further into the surrounding hydrogel. This is consistent with previous studies in agarose hydrogels^23^. In contrast, although collagen type II was evenly distributed at the cell-hydrogel interface, it showed little fibrillar extensions into the hydrogel (Figure 4b). Similarly, collagen type VI - a pericellular collagen around chondrocytes - was restricted to the immediate pericellular interface (Figure 4c). These findings indicate that collagens are spatially heterogenous, and their patterns depend on the specific type. Beyond collagens, proteoglycans are also an important component of the chondrogenic ECM. Thus, we next stained for aggrecan which formed a relatively thick layer with some spatial overlap with the outer, low-intensity region of nECM (Figure 4d). Decorin, another important proteoglycan that regulates collagen fibrillogenesis and matrix micromechanics^24^, remained largely confined to the pericellular nECM region (Figure 4e). These observations are consistent with previous studies suggesting that the pericellular ECM proteins physically displace the hydrogel whereas fibrillar proteins and proteoglycans likely extend into the surrounding hydrogel^4,23^. Furthermore, radial intensity profiles revealed a close spatial overlap between nECM and total collagen, indicating similar distribution patterns. Collagen type II and aggrecan displayed relatively uniform intensity extending from the pericellular region toward the outer matrix. In contrast, collagen type VI and decorin signals were largely restricted to the pericellular zone (Figure 4f). Interestingly, quantifying the projected area of collagen type II showed an increase in ‘high’ hydrogels but a decrease when normalized to the total nECM (Supplementary Figure 4). Other ECM proteins followed a similar trend. These observations highlight that normalizing to nECM may introduce biases. In addition, mask-based area analyses are limited by spatial heterogeneity. Together, these limitations underscore the need for complementary quantification approaches beyond imaging-based analyses.

**Figure 4.**
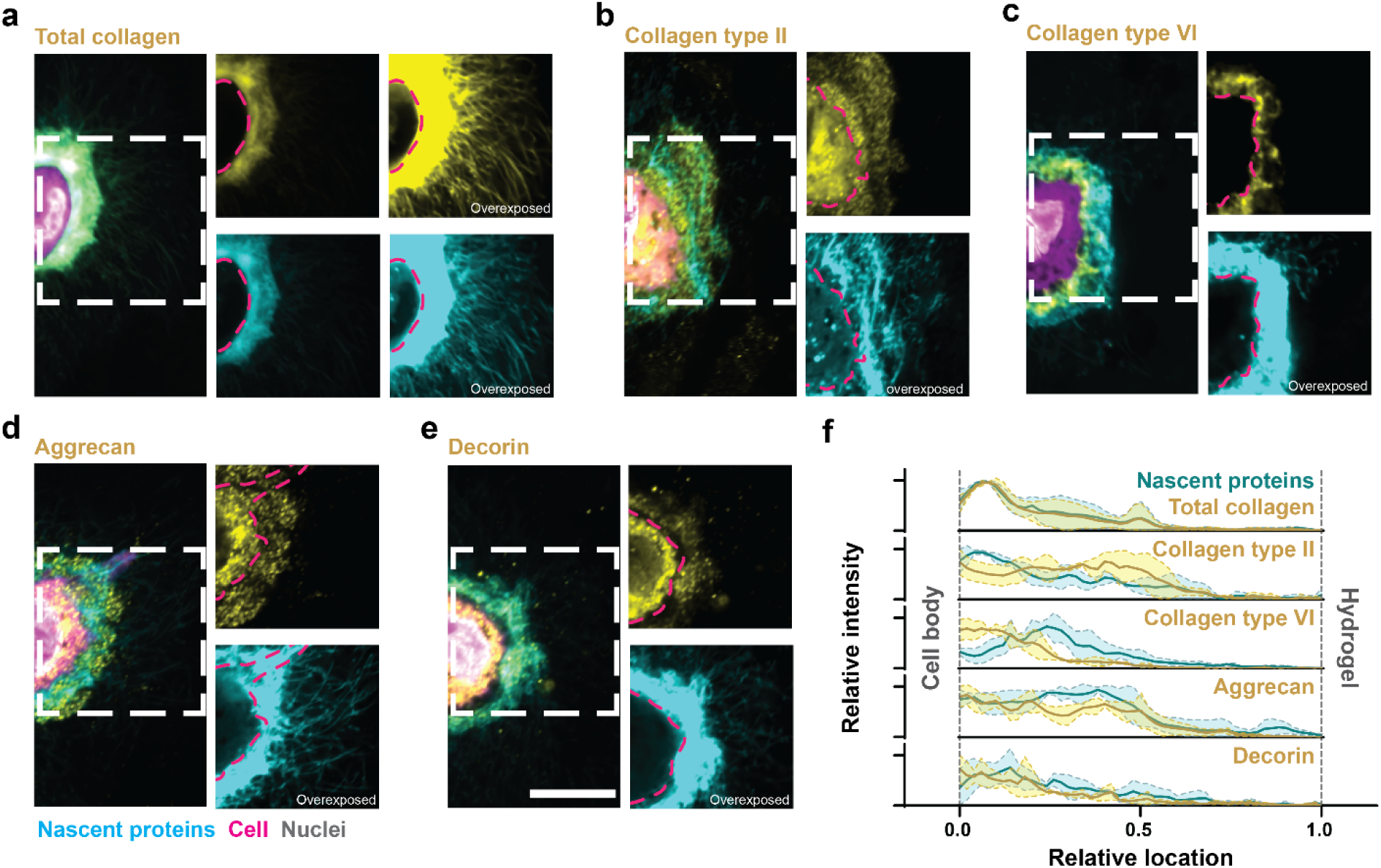
Specific ECM proteins are spatially heterogeneous within the nECM. **a.** Representative fluorescent image of nECM and total collagen deposited by chondrocytes at day 7. **b.** Representative fluorescent image of nECM and collagen type II deposited by chondrocytes at day 7. **c.** Representative fluorescent image of nECM and collagen type VI deposited by chondrocytes at day 7. **d**. Representative fluorescent image of nECM and Aggrecan deposited by chondrocytes at day 7. **e.** Representative fluorescent image of nECM and Decorin deposited by chondrocytes at day 7. **f.** Radial profiles of fluorescence intensities of nECM (cyan) and the specific ECM proteins: total collagen, collagen type II, collagen type VI, aggrecan, and decorin (yellow) at day 7. Lines represent median intensity profiles, shaded areas represent standard deviation (*n* = 3 cells). **a-e.** (scale bar: 10 μm)

### Hydrogel modifications induce differential nECM protein expression

To overcome the limitations of quantification based on immunofluorescence staining, we next performed unbiased bulk proteomics with median normalization. Given the low abundance of nECM proteins within all proteins, we enriched the samples for ECM proteins prior to mass-spectrometry by decellularizing the hydrogels followed by collecting the ECM proteins in high-concentration urea (8M)^6,9^. Next, we used the Matrisome Analyzer^25^ and classified 182 proteins as matrisome components, including the core matrisome (93 total: collagens (23), glycoproteins (54), and proteoglycans (16)) and matrisome-associated proteins (89 total: ECM regulators (44), ECM-affiliated proteins (22), and secreted factors (22) (Figure 5a). Principal component analysis (PCA) for the nECM component abundances revealed clear separation of the proteins within ‘low’ and ‘high’ hydrogels along PC1 (62.7% of the total variance) and PC2 (12.3% of variance, Figure 5b). This tight clustering within each group indicates low technical variability and reproducible measurements across independent experiments. Differential expression was determined using an adjusted *p*-value (*p adj*) threshold of 0.1 (FDR (False Discovery Rate), Benjamini–Hochberg) to maintain sensitivity while controlling for multiple testing in exploratory proteomic datasets. Among the 182 identified matrisome proteins, 55 were significantly different (|fold change| > 2 and *p adj* < 0.1), while 17 showed > 2-fold change only and 26 showed *p adj* < 0.1 only (Figure 5c). Thus, 30% of the identified matrisome proteins showed a significant and more than 2-fold differential expression between ‘low’ and ‘high’ groups. Building upon these results, we next compared specific protein expressions between ‘low’ and ‘high’ groups. Within the core matrisome, several core proteins, including collagen type VI (COL6A3), biglycan (BGN), osteoglycin (OGN) and decorin (DCN, non-significant) were upregulated in ‘low’ compared to ‘high’, suggesting that the nECM is chondrogenic. In contrast, collagen type I (COL1A1), versican (VCAN) and collagen type III (COL3A1) were upregulated in ‘high’ hydrogels, suggesting upregulation of genes linked to fibrotic ECM remodeling (Figure 5d, Supplementary Table 1). Within the matrisome-associated proteins, the differential expression of lysyl oxidases (LOXL2, LOXL3 and LOXL4), matrix metalloproteinases (MMP14), and tissue inhibitor of metalloproteinases (TIMP2 and TIMP3) indicate differences in the regulation of nECM crosslinking, degradation and remodeling between ‘low’ and ‘high’ hydrogels (Figure 4e, Supplementary Table 1). Gene Ontology (GO) analysis performed on the 55 differentially expressed matrisome proteins with the full matrisome (∼1,000 genes)^26^ as the reference background revealed enrichment in articular cartilage development (GO:0061975) and cartilage development (GO:0051216). Enriched protein includes biglycan (BGN), epiphycan (EPYC), osteoglycin (OGN), LOXL2, Matrix Gla Protein (MGP), COL1A1, and COL3A1 (Figure 5f). These enrichments align with gene expression data (Figure 2i) and indicate that the nECM compositional differences in ‘low’ hydrogels are associated with cartilage developmental processes. It further aligns with previous studies showing that low-modification NorHA hydrogels increase cartilage tissue formation in long-term *in vitro* culture^15^.

**Figure 5.**
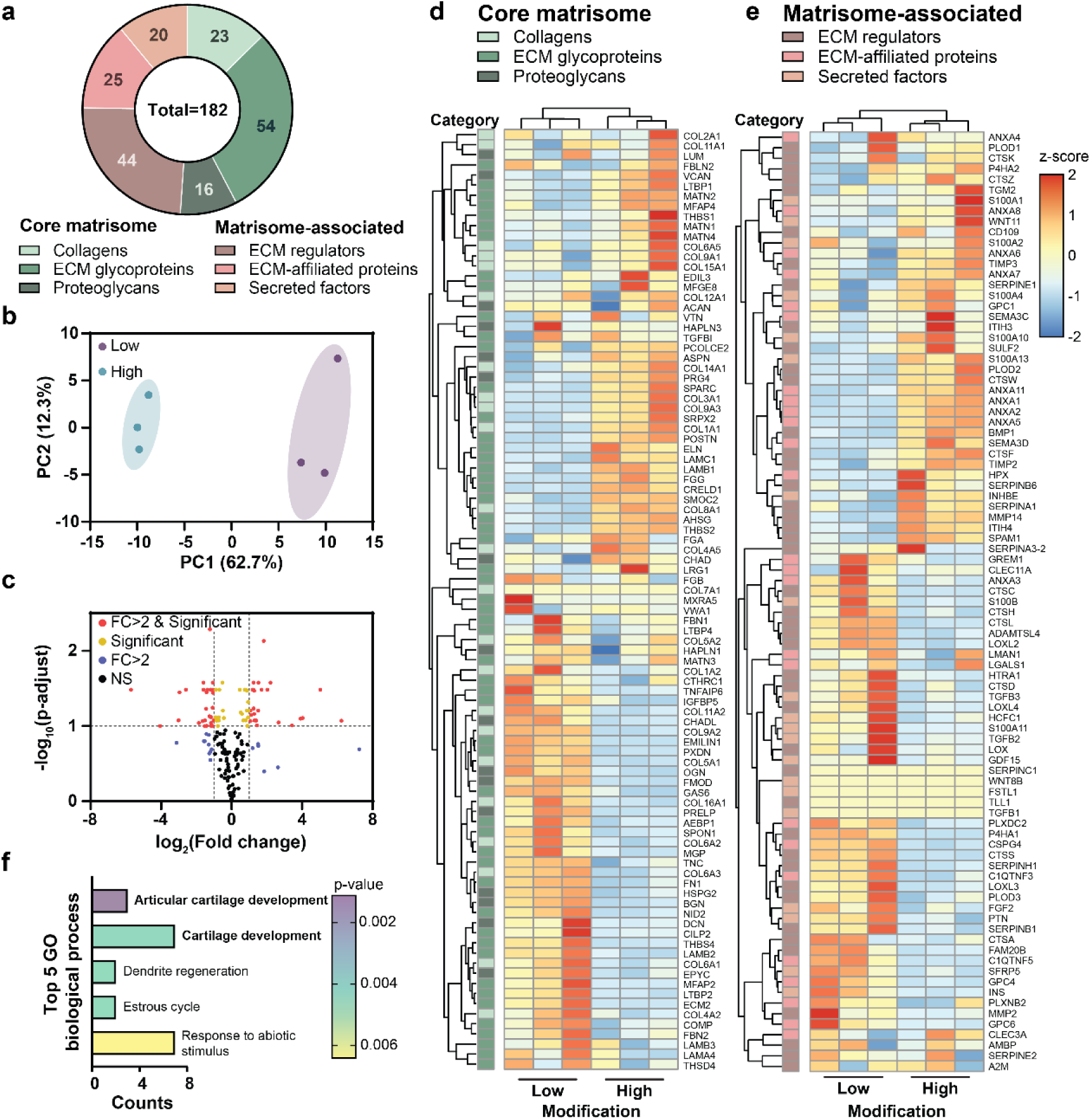
Hydrogel modifications induce differential expression of specific nECM proteins. **a.** Representative pie chart of the distribution of core matrisome deposited by chondrocytes in low and high modification hydrogels including ECM glycoproteins, collagens, and proteoglycans and matrisome- associated proteins including ECM regulators, ECM-affiliated proteins and secreted factors at day 7. **b.** Principal component analysis (PCA) plot of core matrisome and matrisome-associated proteins deposited by chondrocytes in low and high modification hydrogels (*N* = 3 independent experiments). **c.** Volcano plot showing differential expression of core matrisome and matrisome-associated proteins deposited by chondrocytes in high versus low modification hydrogels at day 7. Red dots indicate proteins with differential expression levels and high folds changes (|FC| > 2 and adjusted p < 0.1); yellow dots indicate proteins with only significant by adjusted p (adjusted p < 0.1, |FC| ≤ 2); blue dots indicate high fold-change-only without significance (|FC| > 2, adjusted p ≥ 0.1); black dots indicate non-significant proteins changes (|FC| ≤ 2 and adjusted p ≥ 0.1). (*N* = 3 independent experiments). Heatmaps of **d.** core matrisome protein expression and **e.** matrisome-associated proteins in low and high modification hydrogels at day 7. Rows show individual proteins grouped by hierarchical clustering analysis: columns show the 3 different samples with expression levels (z-score by rows) shown as color intensity from negative z-scores (blue) to positive z-scores (red/yellow). The left color bar denotes protein categories: core matrisome (ECM glycoproteins, collagens, proteoglycans) and matrisome-associated proteins (ECM regulators, ECM-affiliated proteins, and secreted factors). **f.** Top 5 significant Gene Ontology (GO) enrichment pathways with p < 0.05 of biological process, including articular cartilage development, cartilage development, dendrite regeneration, estrous cycle, and response to abiotic stimulus. Counts indicate the number of proteins significantly enriched in each pathway. *N* = number of independent experiments

Taken together, these findings connect changes in nECM deposition and chondrogenic cell fate with distinct changes in the nECM matrisome, suggesting a direct relationship between hydrogel modifications, nECM composition and cell fate.

### Cell-nECMs interactions determine cell fate

Given that hydrogel modifications induce differential expressions of matrisome proteins, we next asked whether this deposited nECM itself contributes to the maintenance of chondrogenic phenotypes. Because cells interact with nECM via integrins, we treated cells with a function-perturbing antibody against integrin β1 (ITGB1i), a major integrin subunit broadly involved in cell adhesion to diverse ECM proteins. Our previous studies further showed that cells deposit nECM within a few hours^4^. Thus, we blocked cell-nECM interactions immediately after embedding and for the entire cell culture period (Figure 6a). Integrin β1 expression was slightly higher in cells cultured in the ‘high’ group from 1-hour post-embedding to Day 7, although ITGB1i treatment had minimal effect on nECM deposition (Supplementary Figure 5). However, inhibition of ITGB1 in ‘low’ hydrogels resulted in an almost 2-fold increase in EdU incorporation with little changes between Ctrl and ITGB1i-treated cells in ‘high’ hydrogels (Figure 6b,c). This suggests that blocking interactions between the cell and chondrogenic nECM in ‘low’ hydrogels leads to similar high cell proliferation in ‘high’ hydrogels. Notably, blocking ITGB1 of cells in ‘low’ hydrogels induced a significant decrease in Sox9 nuclear staining and nuclear-to-cytoplasmic ratio to the same levels as observed in ‘high’ hydrogels (Figure 6d,e). This data shows that cellular interaction with the nECM in ‘low’ hydrogels were providing the pro-chondrogenic signals that are required to maintain cell fate and this goes beyond the initial hydrogel modifications. Importantly, blocking ITGB1 in ‘high’ hydrogels, however, increased Sox9 nuclear staining and nuclear-to-cytoplasmic translocation (Figure 6d, e), suggesting a rescue of cell differentiation. The ability to restore Sox9 cytoplasmic-to-nuclear transition by disrupting cell-nECM interactions suggests that nECM in ‘high’ hydrogels promotes de-differentiation. Together, these findings suggest that the deposited nECM-cell interaction is a critical regulator of cell fate.

#### Outlook

Previous studies have focused on engineering hydrogels with tunable mechanical and biochemical properties to mimic cell-ECM interactions and instruct cell behavior^27^. More recent work has revealed that cells rapidly deposit nECM upon embedding, creating a dynamic interface between cells and the engineered hydrogels^6,23,28^. Yet, the complex interactions between cells, hydrogels and the nECM have not been well studied. Using hydrogels with variable chemical modifications and embedded chondrocytes, we establish a new framework describing a tri-directional interplay among cells, hydrogels, and the deposited nECM (Figure 7). Specifically, we found that low hydrogel modifications led to reduced deposition and accumulation of nECM. However, such nECM was featured in increased chondrogenic proteins. Blocking cellular interactions via integrin β1 inhibition reduced their chondrogenic cell fate. In contrast, high modifications increased nECM accumulation but with less-chondrogenic compositional signatures that promoted proliferation over differentiation. Notably, blocking cellular interactions with ‘high’ nECM rescued chondrogenic cell fate. Thus, our study extends Mina Bissell’s pioneering concept of bidirectional communication between cells and their ECM or engineered hydrogel^2,3^ by revealing the nECM as a third regulatory player that actively governs cell fate decisions in 3D hydrogel systems.

**Figure 6.**
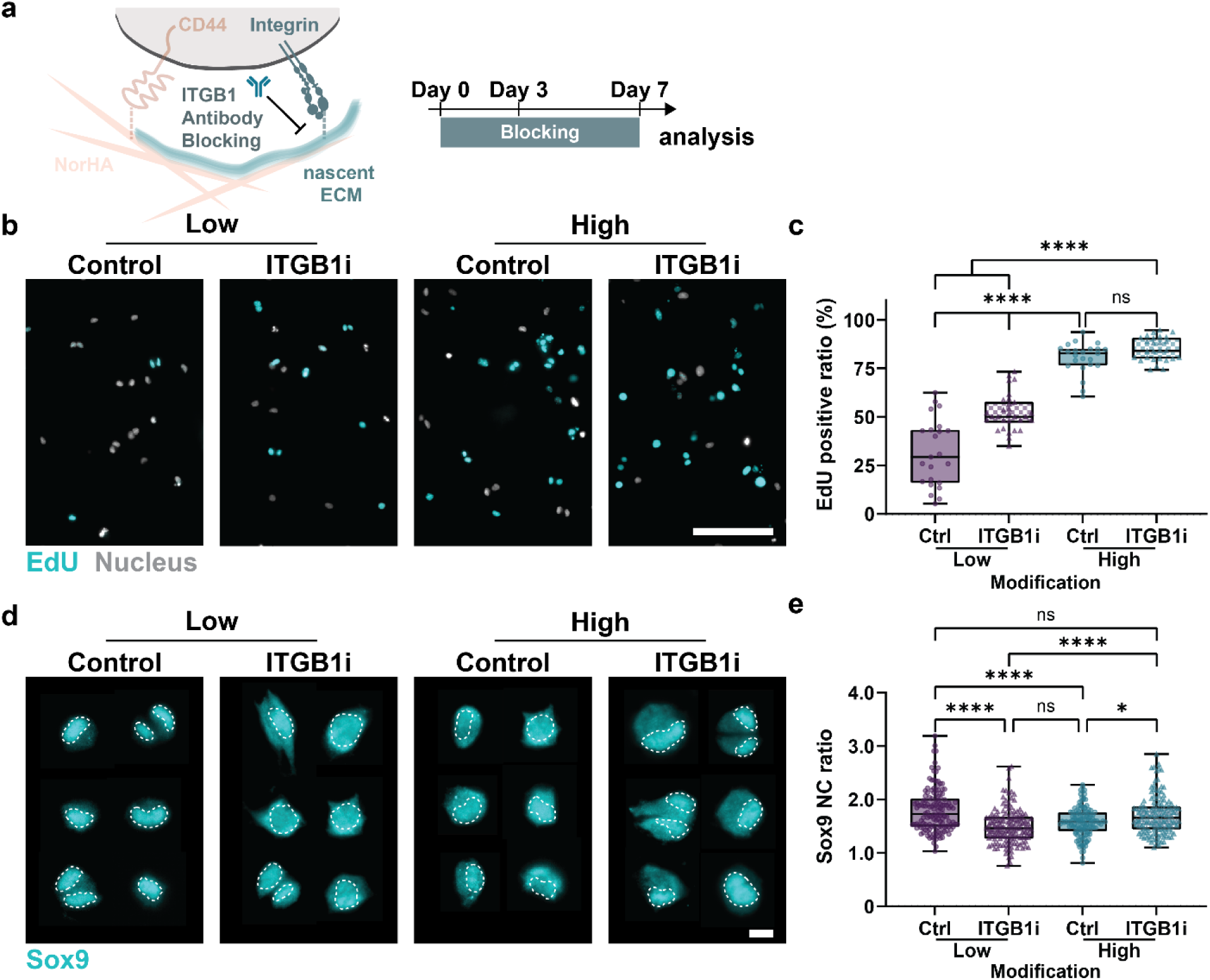
Cell-nECM interaction via integrin β1 controls chondrogenic differentiation. **a.** Timeline and schematic showing the blocking of cell adhesion to nECM using an integrin β1 (ITGB1) function perturbing antibody for 1 hour pre-embedding and for 7 consecutive days**. b.** Representative fluorescent images of the incorporation of EdU in chondrocytes cultured without (Ctrl) and with ITGB1 inhibition (ITGB1i) in high and low modification hydrogels at day 7 (scale bar = 100 μm). **c.** Quantification of the incorporation of EdU in chondrocytes cultured without (Ctrl) and with ITGB1 inhibition (ITGB1i) in high and low modification hydrogels at day 7 (low-Ctrl: *n* = 23 ROIs, *N* = 3; low-ITGB1i: *n* = 30 ROIs, *N* = 3; high-Ctrl: *n* = 23 ROIs, *N* = 3; low-ITGB1i: *n* = 27 ROIs, *N* = 3). **d.** Representative fluorescent images of Sox9 immunofluorescence (dashed line outlines nuclei) of chondrocytes cultured without (Ctrl) and with ITGB1 inhibition (ITGB1i) in low and high modification hydrogels at day 7 (scale bar = 10 μm). **e.** Quantification of Sox9 nucleus-to-cytoplasm (NC) ratio of chondrocytes cultured without (Ctrl) and with ITGB1 inhibition (ITGB1i) in low and high modification hydrogels at day 7 (low-Ctrl: *n* = 138 cells, *N* = 4; low-ITGB1i: *n* = 133 cells, *N* = 4; high-Ctrl: *n* = 124 cells, *N* = 4; high-ITGBi: *n* = 138 cells, *N* = 4). **a-e.** *N* = number of independent experiments ****p < 0.0001, *p < 0.05, error bar = standard deviation, ns: no significant difference by two-way ANOVA with Tukey’s multiple comparisons test.

**Figure 7.**
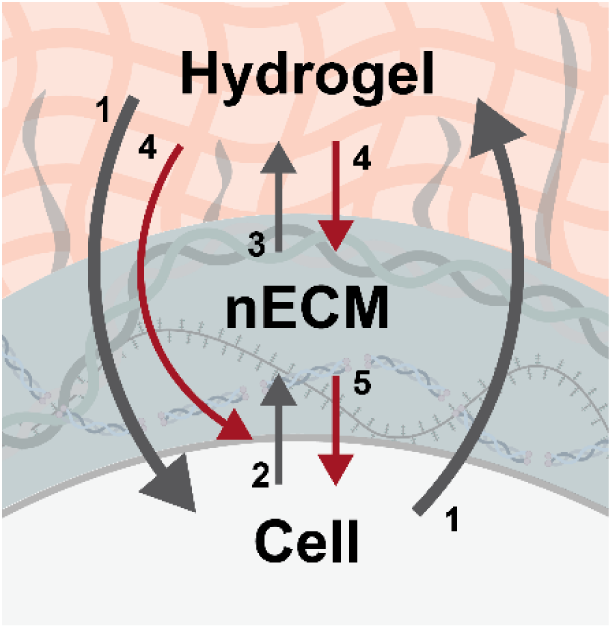
Tri-directional interplay among cells, hydrogels and their nECM. Schematic illustrating previous findings (grey arrows), including the traditional bi-directional interplay between cells and the ECM or engineered hydrogels (**1**, Bissel 1982^2^), and recent work that showed the rapid deposition of nECM after embedding cells into engineered hydrogels (**2**, McLeod 2016; Loebel 2019; Cha 2024^6,23,28^) that interpenetrates into the existing hydrogel (**3**, Loebel 2020^8^). This work (red arrows) demonstrates that hydrogel modifications direct nECM deposition and accumulation (**4**), and induce cell fate decisions (**5**).

In this work, we selected chondrocytes as a well-characterized cell source that responds to hyaluronic acid backbone signaling and is a robust ECM producer^29^. However, additional studies are required to investigate whether this tri-directional framework generalizes across different cell types and hydrogel platforms. Furthermore, several studies have incorporated cell-adhesive moieties and degradable hydrogel crosslinkers which may additionally regulate nECM deposition and interpenetration with the hydrogel^13,30^. Future work incorporating peptides like RGD or collagen-mimetic binding peptides and the use of enzymatically or hydrolytically-degradable peptide crosslinkers will be critical to further expand and validate this tri-directional framework across diverse biomaterial systems.

Taken together, our work demonstrates that hydrogel chemical modifications guide cell fate through the deposition and accumulation of nECM and subsequent cell-nECM interactions. Given the broad applicability and use of engineered hydrogels in mechanobiology and disease modeling, our findings underscore the need to recalibrate hydrogel design parameters and the interpretation of cell fate in 3D hydrogel systems. This tri-directional framework provides a new lens through which to understand cell-biomaterial interactions. It further offers design principles for engineering biomaterials that harness nECM as an instructive intermediary to direct desired cellular outcomes.

## Experimental

### Hydrogel synthesis

NorHA was synthesized via a 4-(4,6-dimethoxy-1,3,5-triazin-2-yl)-4-methylmorpholinium chloride (DMTMM)-mediated aqueous coupling reaction adapted from previously reported methods^16^. Sodium hyaluronate (Lifecore Biomedical, MW ∼68.1 kDa) was dissolved at 1% (w/v) in 0.1 M MES buffer (pH 5.5). DMTMM (TCI, >98%) and 5-norbornene-2-methylamine (Nor, TCI, >98%) were added at defined molar ratios to tune the degree of modification. For low modification, a 1:1:3 HA:DMTMM:Nor ratio was reacted for 24 h. For mid, 1:3:2 was used for 24 h. For high, the same 1:3:2 ratio was used with a second addition of DMTMM and Nor after 24 h, continuing for 48 h total.

Following the reaction, NorHA was precipitated with saturated NaCl and ethanol, then resuspended in Milli-Q water and dialyzed for three days using 6–8 kDa tubing against 0.25 g/L DPBS (Gibco). The final product was frozen, lyophilized, and stored at –20 °C.

To determine the degree of norbornene modification, we performed proton nuclear magnetic resonance (^1^H NMR) spectroscopy. Modified hyaluronic acid products were dissolved in deuterium oxide at 7 mg/mL, and spectra were acquired using a 600 MHz NEO400 spectrometer (Bruker). The modification degree was calculated from the ratio of integrated signal intensities corresponding to backbone protons of hyaluronic acid and vinyl protons of the norbornene groups. (Supplementary Figure 6). Baseline correction and spectral analysis were conducted in MestReNova (v15.1.0, Mestrelab Research).

### Hydrogel formation

Lyophilized NorHA was sterilized using a UV oven (Tool Klean) for 30 minutes. Hydrogels were prepared at 2 wt% by dissolving the sterilized NorHA in buffer (pH 7.5) containing 50 mM HEPES (Gibco) and 1 mg/mL phenol red (Remel). The photoinitiator lithium phenyl-2,4,6-trimethylbenzoylphosphinate (LAP, Arkema Sartomer) was added at a final concentration of 1.70 mM, and dithiothreitol (DTT, Sigma-Aldrich) was included as the crosslinker at designated concentrations ranging from 0.13 to 13 mM The precursor solution was then exposed to UV light (OmniCure, 5 mW/cm²) for 3 minutes to initiate gelation.

### Mechanical characterization

Hydrogels were cast in cylindrical molds (5 mm diameter) and subjected to unconfined compression testing using a Discovery HR-30 Hybrid Rheometer (TA Instruments). Tests were conducted at room temperature with a constant compression rate of 20 µm/s. The Young’s modulus was determined from the linear region of the stress-strain curve, specifically between 10% and 20% strain.

### Cell culture, encapsulation and antibody blocking

Primary chondrocytes were isolated from juvenile bovine articular cartilage as previously described. Briefly, femoral condyles from juvenile bovine knees (6 months old, purchased from Research 87) were dissected to collect articular cartilage. The tissue was digested in 1 mg/mL type II collagenase (Worthington Biochemical) in DPBS for 20 hours at 37 °C in a humidified incubator with 5% CO₂. The resulting cell suspension was filtered through a 70 µm cell strainer to remove debris. Isolated chondrocytes were expanded for one passage on tissue culture-treated dishes at a seeding density of 10,000 cells/cm² in high-glucose DMEM (Gibco) supplemented with 10% fetal bovine serum (FBS, Corning), 1% penicillin–streptomycin (Gibco), and 1% sodium pyruvate (Gibco). Expanded cells were encapsulated in NorHA hydrogels at a density of 5 million cells/mL. Cell-laden hydrogels were cultured for 7 days in chondrogenic-azidohomoalanine (AHA) medium composed of glutamine, L-methionine, and L-cystine-free high-glucose DMEM (Gibco), supplemented with 0.1 µM dexamethasone (Sigma-Aldrich), 4 mM GlutaMAX (Gibco), 0.201 mM L-cystine (Sigma-Aldrich), 50 µM L-methionine(Sigma-Aldrich), 100 µg/mL sodium pyruvate, 1.25 mg/mL bovine serum albumin (BSA, Sigma-Aldrich), 0.1% ITS+ premix (Gibco), 50 µg/mL ascorbate-2-phosphate(Sigma-Aldrich), 40 µg/mL L-proline (Sigma-Aldrich), and 1% penicillin–streptomycin–amphotericin. Media was further supplemented with 10 ng/mL Transforming Growth Factor Beta (TGFβ)-3 (R&D system) and 50 µM L-AHA (Vector Lab).

For perturbation studies, media was supplemented with either an anti-CD44 antibody (DSHB, H4C4, 2.5 µg/mL, day 0-3) or anti-integrin β1 antibody (anti-ITGB1, DSHB, AIIB2, 2.5 µg/mL, day 0-7). For CD44 blocking studies cells were additionally incubated in DPBS with anti-CD44 for 1 hour prior to encapsulation^31^.

### Cell proliferation assays

For divided cell quantification, cells were stained with Hoechst 33342 (Invitrogen) and CellMask Deep Red (Invitrogen) and fixated with 4% paraformaldehyde. Cells were classified as “divided” when either multiple nuclei were observed within a single continuous membrane boundary or two closely apposed cells remained physically connected, consistent with recently divided daughter cells. For EdU incorporation assays, 5 µM EdU was added to chondrogenic AHA media, and incorporated EdU was labeled after fixation using the Click-&-Go® Cell Reaction Buffer Kit (Vector Labs, CCT-1263) and AZDye 647 Azide (Click Chemistry Tools, 1482-1) via copper-catalyzed click chemistry. Imaging was performed blindly by selecting random regions using a Leica THUNDER microscope (40× objective), acquiring Z-stacks spanning 100 µm starting from 50 µm below the hydrogel surface. The divided cell ratio was calculated as the number of divided cells divided by the total number of cells, while EdU incorporation was determined by the ratio of EdU-positive nuclei to total nuclei.

### Nascent matrix labeling and quantification

Nascent matrix metabolic labeling was performed following a previously established protocol^9^. Briefly, cells were stained with 30 µM AZDye 488 DBCO (Vector Labs), Hoechst, and CellMask, followed by fixation and washing overnight. Imaging was performed blindly by selecting random regions using a Leica THUNDER microscope (40× objective), acquiring Z-stacks spanning 100 µm starting from 50 µm below the hydrogel surface. Individual cells were cropped manually, and unbiased image analysis was conducted using an ImageJ macro. Otsu auto-thresholding was applied to both the nECM and CellMask channels. To isolate the extracellular region of nECM, the CellMask signal was subtracted from the nECM channel. The average thickness of nECM of individual cells was calculated using the BoneJ plugin, while volumes of the cell and nECM were quantified using the 3D Viewer plugin.

### Gene expression

Total mRNA was isolated from encapsulated cells. Briefly, hydrogels were degraded using 2 mg/mL hyaluronidase (Sigma-Aldrich) by 30 min incubation at 37°C, followed by RNA extraction using the TRIzol-chloroform method. RNA quality was confirmed by NanoDrop One spectrophotometer (Thermo Fisher Scientific), and RNA concentration was measured using the Qubit™ RNA High Sensitivity (HS) Assay Kit (Invitrogen, Q32852). cDNA synthesis was performed using the High-Capacity cDNA Reverse Transcription Kit (Applied Biosystems, Invitrogen, 4368814) according to the manufacturer’s protocol. Quantitative PCR (qPCR) was conducted using PowerUp™ SYBR™ Green Master Mix (Invitrogen, A25742) on an Applied Biosystems QuantStudio 3 Real-Time PCR System, with three technical replicates per sample.

Gene expression levels were calculated using the ΔΔCt method, with Ribosomal Protein S18 (RPS18) as the reference gene. Expression levels were normalized to the ‘low’ group. In addition to individual gene expression analysis, COL2A1/COL1A1 and ACAN/VCAN expression ratios were also calculated to assess chondrogenic differentiation and matrix composition^32–35^.

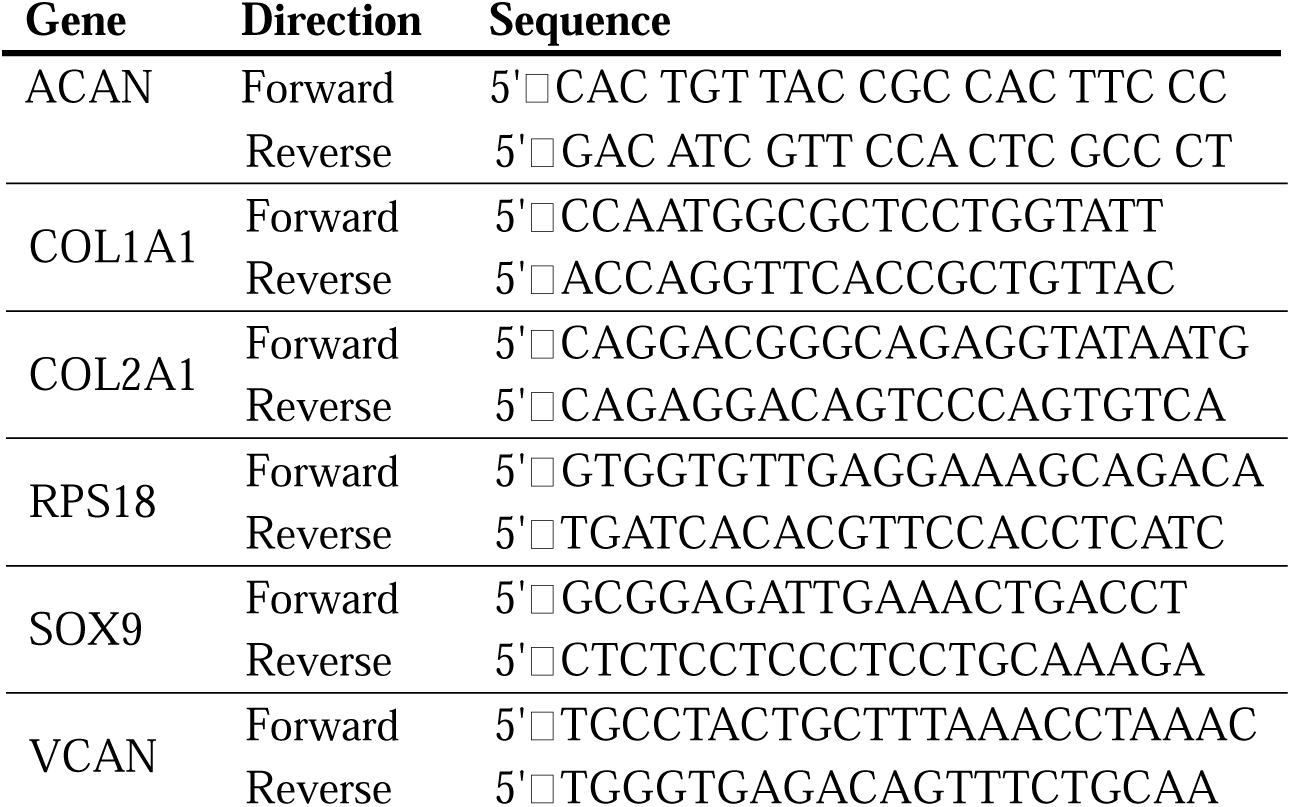

### Immunofluorescence staining, imaging and analyses

Samples were fixated and cryoprotected by sequential incubation in 30% sucrose overnight, followed by 2 hours in a 1:1 mixture of sucrose and OCT compound, and 1 hour in 100% OCT. Samples were embedded in OCT and placed into a room-temperature 2-methylbutane bath (Thermo Fisher Scientific), which was then snap-frozen by immersion in liquid nitrogen to minimize ice crystal formation. Cryosectioning was performed at −23 °C using a Leica CM3050S cryostat to obtain 10 µm thick slices. Slides were baked at 37 °C for 30 minutes before storage at −20 °C. Cryosections were blocked in 5% (w/v) BSA for 30 minutes, incubated with primary antibodies diluted in 2% BSA overnight at 4 °C, and then incubated with secondary antibodies in 2% BSA for 1 hour at room temperature. DAPI (Invitrogen, 62248, 1:1000) and HCS cytoplasm CellMask Deep Red (Invitrogen, H32721, 1:5000) were included during secondary antibody incubation. Slides were mounted using SlowFade Diamond antifade mountant with DAPI (Invitrogen, S36968). Primary antibodies included SOX9 (NovusBio, NBP2-24659, 1:100), Collagen II (DSHB, II-II6B3, 1:100), Collagen VI (Biosynth, 70R-CR009X, 1:100), Decorin (Kerafast, ENH077-FP, 1:100), and Aggrecan (Abcam, ab3778, 1:50). Secondary antibodies included goat anti-mouse IgG Alexa Fluor 568 (Invitrogen, A11031, 1:200) and goat anti-rabbit IgG Alexa Fluor 568 (Invitrogen, A11011, 1:200).

Total collagen content was visualized using a Cy3-conjugated collagen hybridization peptide (CHP, 3 Helix, RED60^36^). Slides were first subjected to heat-induced collagen denaturation by steaming at 95 °C for 25 minutes. Staining was then performed following the manufacturer’s instructions. Briefly, the CHP solution was rapidly heated, immediately cooled on ice, and applied to the slides. Samples were incubated at 4 °C overnight to allow peptide binding to denatured collagen strands. Imaging was performed using either a Nikon Eclipse Ti2-E microscope (60× objective) or a Leica THUNDER microscope (40× objective). The regions of interest were chosen blindly and mages acquired at the midplane of the cell body.

Calculation of SOX9 nucleus-to-cytoplasm ratio was calculated for each individual cell. The DAPI channel was used to generate a nuclear mask, while the cytoplasmic region was defined by subtracting the nuclear mask from the CellMask channel. The mean fluorescence intensity of SOX9 within each region was measured, and the nuclear-to-cytoplasmic ratio was calculated by dividing the nuclear intensity by the cytoplasmic intensity.

nECM thickness analyses were performed using fluorescence intensity profiles by drawing a line extending from the cell edge to the farthest detectable nECM signal within the hydrogel. Fluorescence intensity was recorded along this line. Both position and intensity values were normalized, with position defined as 0 at the cell surface and 1 at the outermost nECM boundary, and intensity scaled from 0 (minimum) to 1 (maximum). Intensity profiles from three representative cells were analyzed to illustrate the trend. Averaged data were then used to generate plots showing the mean intensity and standard deviation. For nECM composition analysis, protein-specific and nECM signals were processed by first excluding intracellular regions using the whole cell body mask. The remaining extracellular signal was identified using Otsu auto-thresholding. The relative ratio was calculated as the area of protein-positive signal divided by the area of total nECM signal.

### Proteomics and matrisome analysis

Protein isolation and nECM enrichment were performed based on a previously described protocol. Briefly, hydrogels were snap-frozen in liquid nitrogen and subsequently decellularized using 1.5 M potassium chloride (KCl, Sigma-Aldrich) with 0.1% Triton X-100 (Sigma-Aldrich) in 50 mM Tris-HCl (Sigma-Aldrich) buffer at pH 8.0 on ice for 6 hours in the presence of a protease inhibitor cocktail (cOmplete™, Roche). Residual hydrogel and DNA were enzymatically digested overnight at 37 °C using 0.5mg/mL hyaluronidase and DNase (GLPBIO), with continued protease inhibition.

After discarding the supernatant, pelleted protein was solubilized in 8 M urea (Thermo Fisher Scientific) prepared in 50 mM ammonium bicarbonate buffer (Sigma-Aldrich), followed by acetone precipitation at a 1:4 ratio and overnight incubation at 4 °C. Protein concentration was determined using the BCA assay (Thermo Fisher Scientific, A55864), and 40 µg of protein from each sample was used for downstream analysis.

Samples were labeled with the TMTsixplex™ Isobaric Label Reagent Set (Thermo Fisher Scientific) and submitted to the University of Michigan Proteomics Core for tandem mass tag (TMT)-based mass spectrometry on a fee-for-service basis. Protein identification and quantification were performed using Proteome Discoverer 3.0. Data were searched against the Bos taurus UniProt database (sp_tr_canonical, TaxID 9913, v2023-06-28), allowing dynamic modifications including methionine oxidation (+15.995 Da), deamidation (+0.984 Da, N/Q), and methionine-to-azidohomoalanine substitution (−4.986 Da, M). Trypsin was used as the digestion enzyme, and results were filtered at 1% false discovery rate (FDR) for high-confidence peptide and protein identification.

Annotation of matrisome proteins, including collagens, glycoproteins, proteoglycans, ECM-affiliated proteins, ECM regulators, and secreted factors, was performed using Matrisome AnalyzeR, based on the bovine matrisome classification and the Matrisome Project database^25,26,37^. Within each sample, matrisome protein abundances were normalized by dividing by the mean abundance of all detected matrisome proteins. Differential expressions were assessed by calculating Z-scores of each protein across samples.Hierarchical clustering was performed on the normalized matrisome subset to generate heatmaps, while principal component analysis (PCA) and volcano plots were constructed based on differential expression data. Gene Ontology (GO) over-representation analysis was conducted using the WEB-based GEne SeT AnaLysis Toolkit (WebGestalt, https://www.webgestalt.org/). Differentially expressed matrisome proteins (fold change > 2 and *p-adjust* < 0.1) were used as the input list, with the full matrisome database serving as the reference background. The analysis was performed using the *Bos taurus* genome as the organism of interest and “Biological Process” as the functional category. Statistical significance was determined by Fisher’s exact test, and multiple testing correction was applied using the Benjamini-Hochberg (BH) method.

### Flow cytometry of integrin β1

Cell-laden hydrogels were fixed in 4% paraformaldehyde for 30 minutes and digested with 2 mg/mL hyaluronidase overnight at 37 °C to release encapsulated cells. The resulting cell suspension was treated with DNase I (Qiagen) and washed in FACS buffer containing PBS without calcium and magnesium, 0.5% (w/v) BSA (Sigma-Aldrich), 5 mM EDTA (Sigma-Aldrich), and 0.1% sodium azide(Sigma-Aldrich). Cells were blocked in FACS buffer for 30 minutes at room temperature, incubated with a primary antibody against integrin β1 (Invitrogen, PA5-78028, 1:200) for 30 minutes at room temperature, and subsequently incubated with a secondary antibody, goat anti-rabbit IgG Alexa Fluor 647 (Invitrogen, A21244, 1:200) and Hoechst 33342 (Invitrogen), for 30 minutes at room temperature. Stained cells were analyzed using a Attune NXT 4 Laser Flow Cytometer (Thermo Fisher Scientific), and data were processed with FlowJo (v10.8, BD Life Science).

### Statistical analysis

Statistical analyses were performed using GraphPad Prism (v10.6) and RStudio (v2025.05.1). Unpaired two-tailed Student’s *t*-tests with Welch’s correction were applied for comparisons between two groups, and one-way or two-way ANOVA was used for comparisons among multiple groups. Data visualization was performed using GraphPad Prism. In all figures, *n* represents individual cells or regions of interest (ROIs), and *N* represents independent experiments. Statistical analyses were performed using *n* as the unit of analysis.Experiments were conducted using primary cells from three independent donors, with independent experiments performed using cells derived from these donors across different experimental conditions.

## Data availability

The data supporting the findings of this study are available in the main article, the Supplementary Information, and the Source Data file. Raw data files are available from the corresponding author upon request.

## Acknowledgements

This work was partially supported by funding from the NIH (R00-HL151670 and R35GM157063 to C.L., R01AR082348 to M.K.), the American Lung Association (IA-939940 to C.L.), and the David and Lucile Packard Foundation (to C.L.).

## Author contributions

J.Y. Liu conceived and designed the study, performed all experiments, analyzed the data, prepared the figures, and drafted the manuscript. E.M. Plaster, M. Fan, D. Ahmed, A. Roy, P. Duran, and P. Panovich contributed to experimental execution and data acquisition. A.S. Piotrowski-Daspit and C.A. Aguilar provided experimental assistance and resources. M.L. Killian contributed intellectual input. C. Loebel supervised the project, provided conceptual guidance, secured funding, and critically revised the manuscript. M.L. Killian and C. Loebel jointly contributed to funding acquisition. All authors contributed to the review and editing of the manuscript and approved the final version for submission.

