## Supplement for "Nascent extracellular matrix converts biomaterial cues into cell fate decisions"

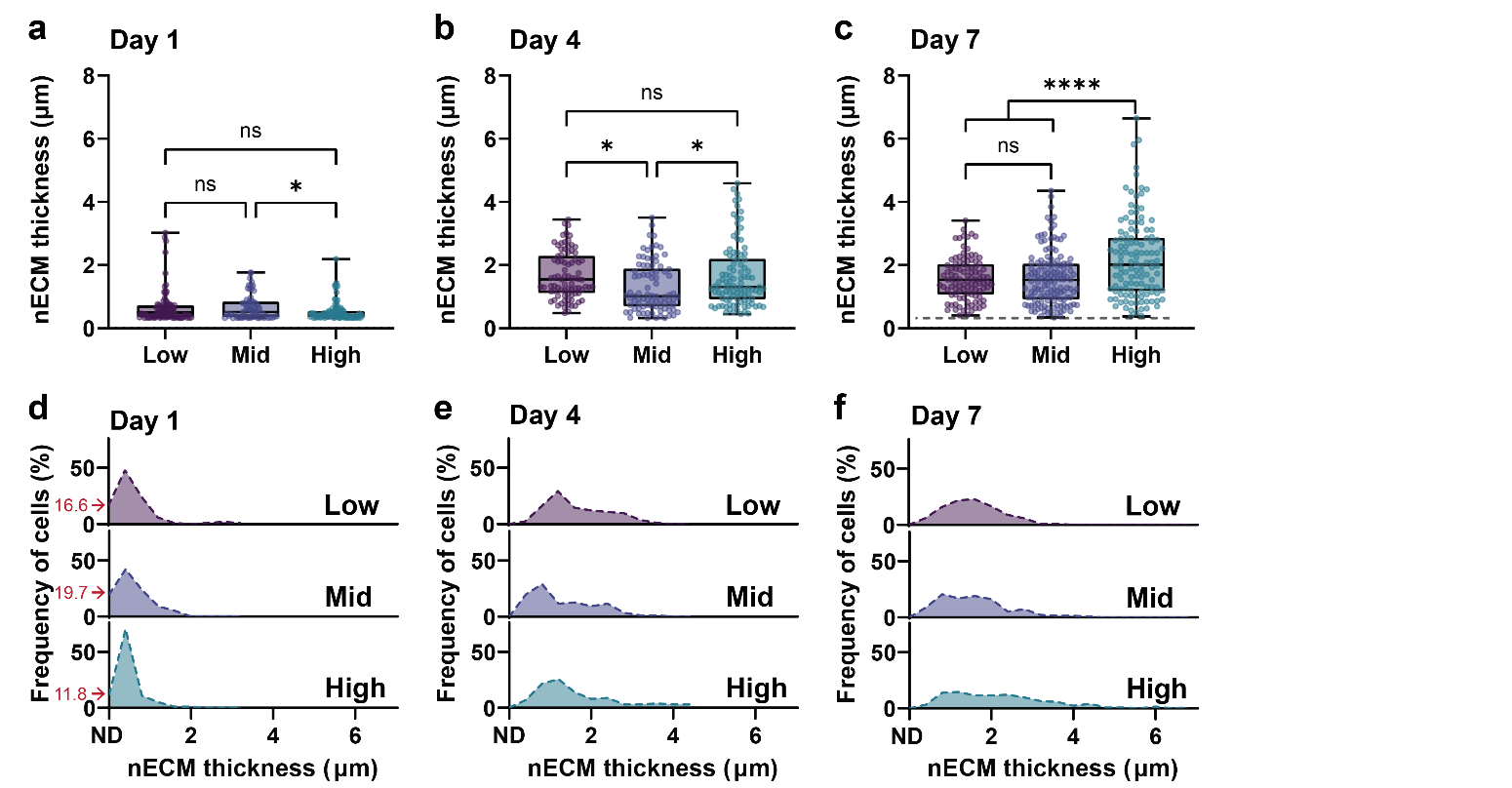

**Figure S1. nECM deposition quantification and heterogeneity over the culture period.**

**a-c.** Quantification of average nECM thickness of chondrocytes cultured in low, mid and high modification hydrogels at day 1, 4, 7 (Day 1: low *n* = 85, *N* = 2; mid *n* = 86, *N* = 2; high *n* = 89, *N* = 2; Day 4: low *n* = 82, *N* = 2; mid n = 86, *N* = 2; high *n* = 102, *N* = 2; Day 7: low *n* = 101, *N* = 3; mid *n* = 136, *N* = 3; high *n* = 122, *N* = 3). **d-f.** Histograms of nECM thickness distribution of chondrocytes cultured in low, mid and high modification hydrogels at day 1, 4, 7 (bin width = 0.4 μm. Red arrows indicate percentages of cells with non-detectable (ND) nECM, Day 1: low *n* = 85, *N* = 2; mid *n* = 86, *N* = 2; high *n* = 89, *N* = 2, Day 4: low *n* = 82, *N* = 2; mid *n* = 86, *N* = 2; high *n* = 102, *N* = 2, Day 7: low *n* = 101, *N* = 3; mid *n* = 136, *N* = 3; high *n* = 122, *N* = 3). **a-f.** *N* = number of independent experiments, error bar = standard deviation, ****p < 0.0001, **p < 0.01, *p < 0.05, ns: not significant by one-way ANOVA with Tukey’s multiple comparisons test.

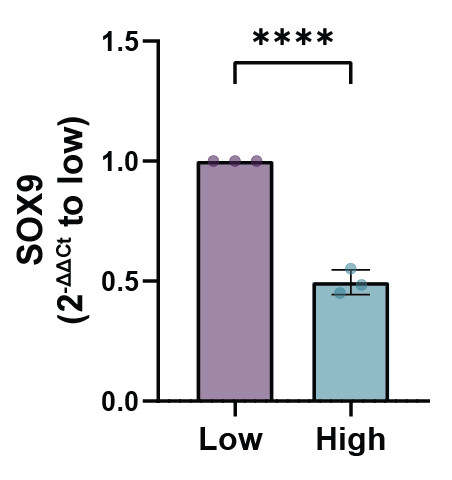

**Figure S2. SOX9 gene expression.** SOX9 gene expression measured by qPCR at day 7, normalized to S18 housekeeping gene using the ΔΔCt method. Data are expressed as fold change relative to low modification hydrogels (set as 1.0). (*N* = 3). *N* = number of independent experiments, error bar = standard deviation, ****p < 0.0001 by Student’s *t-test*.

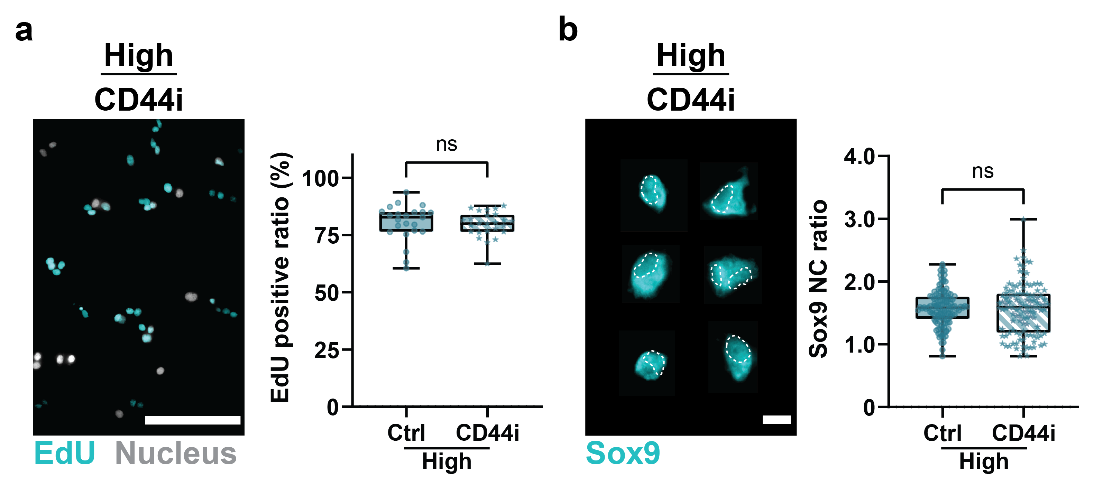

**Figure S3. CD44 blocking in ‘high’ hydrogels.**  **a.** Representative fluorescent images and quantification of the incorporation of EdU in chondrocytes cultured without (Ctrl) and with CD44 inhibition (CD44i) in high modification hydrogels at day 7. (scale bar = 100μm, high-Ctrl: *n* = 23 ROIs, *N* = 3; high-CD44: *n* = 25 ROIs, *N* = 2) **b.** Representative fluorescent images (dashed line outlines nuclei) and quantification of Sox9 nucleus-to-cytoplasm (NC) ratio of chondrocytes cultured without (Ctrl) and with CD44 inhibition (CD44i) in low modification hydrogels at day 7. (scale bar = 10μm, high-Ctrl: *n* =124 cells, *N* = 4; high-CD44: *n* = 117 cells, *N* = 4). **a-b.** *N* = number of independent experiments, error bar = standard deviation, ns: no significant difference by Student’s *t-test*.

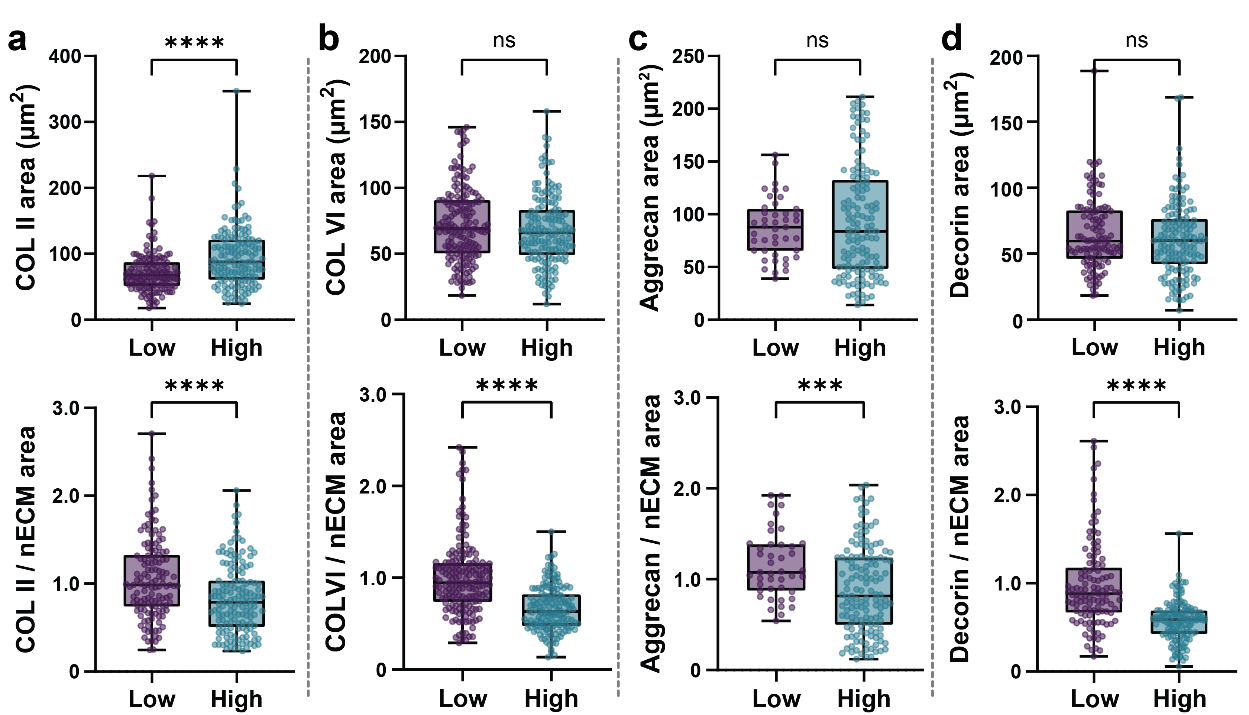

**Figure S4 nECM and specific ECM protein quantifications of mid-plane regions at day 7. a.** Quantification of collagen type II area without normalization and normalized to total nECM area per cell (low: *n* = 132 cells, *N* = 2; High: *n* = 147 cells, *N* = 2). **b.** Quantification of collagen type VI area without normalization and normalized to total nECM area (Low: *n* = 167 cells, *N* = 2; High: *n* = 156 cells, *N* = 2). **c.** Quantification of aggrecan area without normalization and normalized to total nECM area (low: *n* = 43 cells, *N* = 1; High: *n* = 139 cells, *N* = 2). **d.** Quantification of decorin area without normalization and normalized to total nECM area (low: *n* = 114 cells, *N* = 2; High: *n* = 142 cells, *N* = 2). **a-d.** *N* = number of independent experiments, error bar = standard deviation, ****p < 0.0001, ***p < 0.001, ns: no significant difference by Student’s *t-test*.

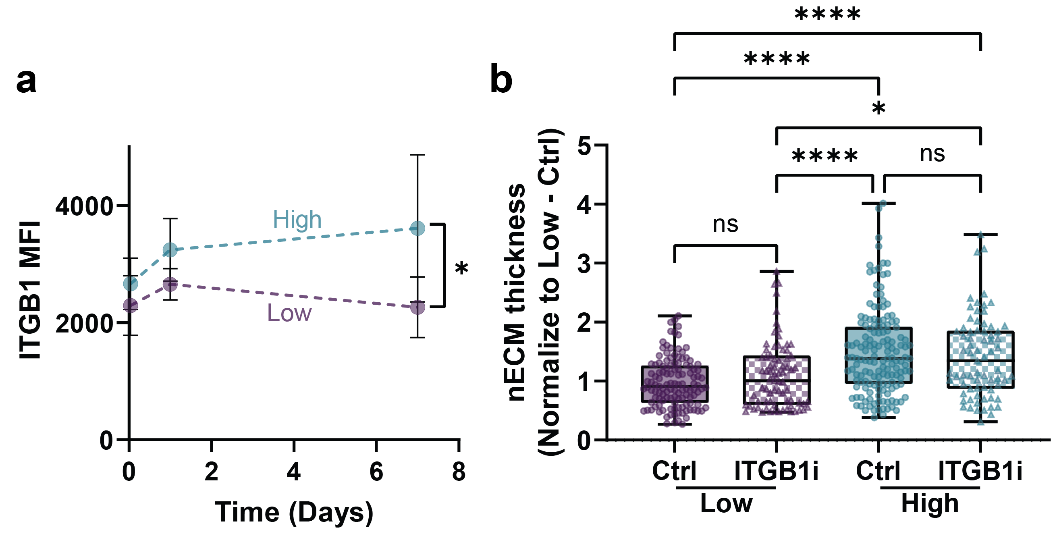

**Figure S5. Quantification of integrin β1 expression and nascent ECM deposition in ITGB1i-treated chondrocytes. a.** Mean fluorescence intensity (MFI) of integrin β1 over 1 hour, day 1, and day 7 following hydrogel encapsulation. **b.** Quantification of nECM thickness of chondrocytes cultured in ‘low’ and ‘high’ hydrogels and treated without or with ITGB1i at day 7. Data were normalized to nECM thickness of ‘low’ Ctrl hydrogels (*n* = 123 cells, *N* = 4; low-ITGB1i: *n* = 75 cells, *N* = 2; high-Ctrl: *n* = 150 cells, *N* = 4; high-ITGB1i: *n* = 72 cells; *N* = 2). *N* = number of independent experiments, error bar = standard deviation, **a:** *p<0.05 by two-way ANOVA with šídák's multiple comparisons test. **b:** ****p < 0.0001, *p<0.05, ns: no significant difference by one-way ANOVA with Tukey’s multiple comparisons test.

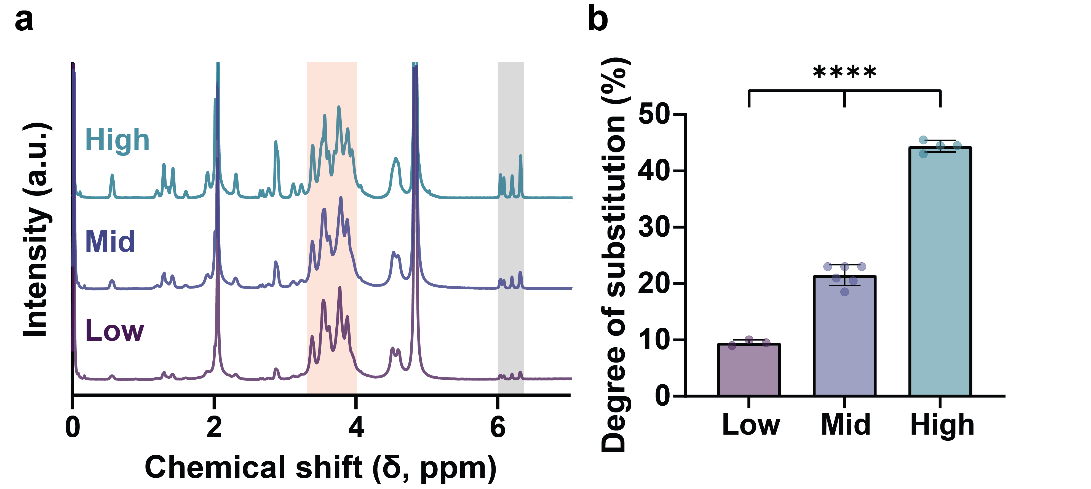

**Figure S6. Chemical characterization of norbornene-modified hyaluronic acid (NorHA) polymer.**

**a.** Representative ^1^H NMR spectrum of NorHA with ‘low’, ‘mid’, and ‘high’ modification. Orange-shaded regions indicate sugar ring protons of HA, while grey-shaded regions correspond to vinyl proton peaks of norbornene groups^28^. **b.** Quantification of the degree of substitution (DS) for the NorHA batches used in this study (‘low’: *n* = 3, *N* = 3; mid’: *n* = 6, *N* = 6, ‘high’: *n* = 5, *N* = 5). *N* = number of independent experiments, error bar = standard deviation, **b:** ****p < 0.0001 by one-way ANOVA with Tukey’s multiple comparisons test.
